# Developmental Normalization of Phenomics Data Generated by High Throughput Plant Phenotyping Systems

**DOI:** 10.1101/2020.05.17.100917

**Authors:** Diego Lozano-Claros, Xiangxiang Meng, Eddie Custovic, Guang Deng, Oliver Berkowitz, James Whelan, Mathew G. Lewsey

## Abstract

**Background:** Sowing time is commonly used as the temporal reference for *Arabidopsis thaliana* (Arabidopsis) experiments in high throughput plant phenotyping (HTPP) systems. This relies on the assumption that germination and seedling establishment are uniform across the population. However, individual seeds have different development trajectories even under uniform environmental conditions. This leads to increased variance in quantitative phenotyping approaches. We developed the Digital Adjustment of Plant Development (DAPD) normalization method. It normalizes time-series HTPP measurements by reference to an early developmental stage and in an automated manner. The timeline of each measurement series is shifted to a reference time. The normalization is determined by cross-correlation at multiple time points of the time-series measurements, which may include rosette area, leaf size, and number.

**Results:** The DAPD method improved the accuracy of phenotyping measurements by decreasing the statistical dispersion of quantitative traits across a time-series. We applied DAPD to evaluate the relative growth rate in *A. thaliana* plants and demonstrated that it improves uniformity in measurements, permitting a more informative comparison between individuals. Application of DAPD decreased variance of phenotyping measurements by up to 2.5 times compared to sowing-time normalization. The DAPD method also identified more outliers than any other central tendency technique applied to the non-normalized dataset.

## Background

Sowing-time is often taken as the initial time point for measuring plant phenotypic traits in HTPP systems [1]. Traits of a number of plants are typically measured between two defined timepoints, chosen relative to the time of sowing. However, the germination of individual seeds in a genetically identical population is normally distributed, even under uniform conditions [2]. Differences in the timing of germination and seed establishment increase the dispersion of time-series measurements because individual plants will be at different developmental stages at any given time point. This may create difficulty in drawing reliable conclusions from the data.

Data normalization is often applied to analyze datasets that have high dispersion, but many methods are not suitable for time-series plant phenotyping data. Traditional methods such as the z-score, min-max, and decimal scaling are not appropriate because of statistical parameters such as mean and standard deviation change over time. Time-series normalization methods can numerically fit the time-series measurements to a single timeline and reduce dispersion. However, these methods do not take into consideration developmental information nor the effect of the allometric scaling of growth; individual seedlings can have similar trait values but maybe at different developmental stages.

Plants progress through specific developmental stages that are consistent between genetically identical individuals grown in uniform conditions. For example, the adjusted BASF, Bayer, Ciba-Geigy (BBCH) scale describes *A. thaliana* developmental stages using seed germination, leaf development, rosette growth, inflorescence emergence, flower production, silique ripening and senescence as significant markers [3]. Considering the availability of clear developmental stage scales, development normalization could be an appropriate method to normalize HTPP data. Developmental normalization would arrange time-series plant phenotype measurements based upon plants being at similar developmental stages. However, developmental normalization has not previously been implemented in an automated manner suitable for image-based HTPP datasets, likely due to the technical difficulty [4].

Image-based plant phenotyping systems are now widely used [5]. However, image processing remains one of the most considerable difficulties for these systems, especially image segmentation of shoots and leaves. Performance depends heavily on the complexity of images, which frequently include interference, light reflection, leaf overlap, and foreign objects that must be removed (for example, pots, and soil) [6]. Furthermore, the identification of multiple leaves at the same time (multi-instance segmentation) is difficult due to their similarity in shape and appearance [4]. Without algorithms that extract accurate measurements, it is more complicated to scale the principal environmental variables influencing the phenotype and underlying physiological processes [7].

Several approaches have been developed to address the challenge of image segmentation. They can be categorized into four groups; shape analysis, watershed-based, machine learning, and graph-based. Shape analysis algorithms rely on assumptions regarding plant geometrical features and structure, but they may fail when encountering new data, which limits their applicability [8], [9]. Watershed-based methods consider a grey image as a topographic surface produced by its intensity gradients, where light pixels are represented as high-intensity values and dark pixels as low-intensity values [4]. However, the performance of watershed algorithms is compromised due to over-segmentation when leaves overlap. Machine learning segmentation approaches can be unsupervised or supervised. Unsupervised learning algorithms are mainly used for pixel clustering. They identify individual leaves by grouping pixels which share a similar feature pattern such as color, texture, and others. Supervised learning algorithms analyze and compare the input plant images with annotated images or labels [10]. Graph-based methods segment individual leaves by applying graph-based noise removal and region growing techniques [8].

We developed DAPD, which combines time-series trait measurement with normalization by developmental stage in an automated manner. DAPD synchronizes the start-point of timelines for all plants in the population by their number of leaves. To achieve this, we also developed a new leaf segmentation algorithm that utilizes both shape analysis and supervised machine learning algorithms. This combined approach overcomes the limitations of individual segmentation methods. We applied DAPD to evaluate the rosette area in *A. thaliana* and demonstrated that it improved uniformity in measurements, enabling a more informative comparison between individuals. The algorithm allows users to select and define the starting leaf number relevant to their experiment. Our code is available for reuse at https://github.com/diloc/DAPD_Normalization.git.

## Results

### DAPD time normalization of plant phenomics data

Our major aim was to produce an automated method for the developmental stage synchronization of HTPP data. To achieve this, we developed DAPD, which synchronizes shoot phenotypic measurements of multiple Arabidopsis plants by normalizing time-series measurements to a reference time point (i.e., a day). DAPD does so by identifying the highest cross-correlation score between leaf number and day-after-sowing (DAS) of seedlings of the same genotype. First, seedlings were grouped by genotype, then their leaf number was assessed by applying the DAPD-Leaf Counting algorithm. Next, a daily leaf number was associated with a specific day in the DAS timeline, and the degree of similarity between them was calculated by cross-correlation. The highest score indicates the day where the leaf number of time-series are best aligned. We intended that the application of DAPD would decrease the dispersion of time-series phenotype measurements.

We tested DAPD on two independent experiments comprised of many *A. thaliana* ecotype Col-0 and Cvi replicates. In the first experiment, we used 355 individuals of Col-0 only and, in the second experiment, 140 individuals of each ecotype. Replicate plants of the same ecotype within experiments were at different developmental stages, as assessed by leaf number, despite being sown at the same time and having been grown from seeds of plants cultivated under conditions that would not result in seed dormancy (Figure 1).

**Figure 1.**
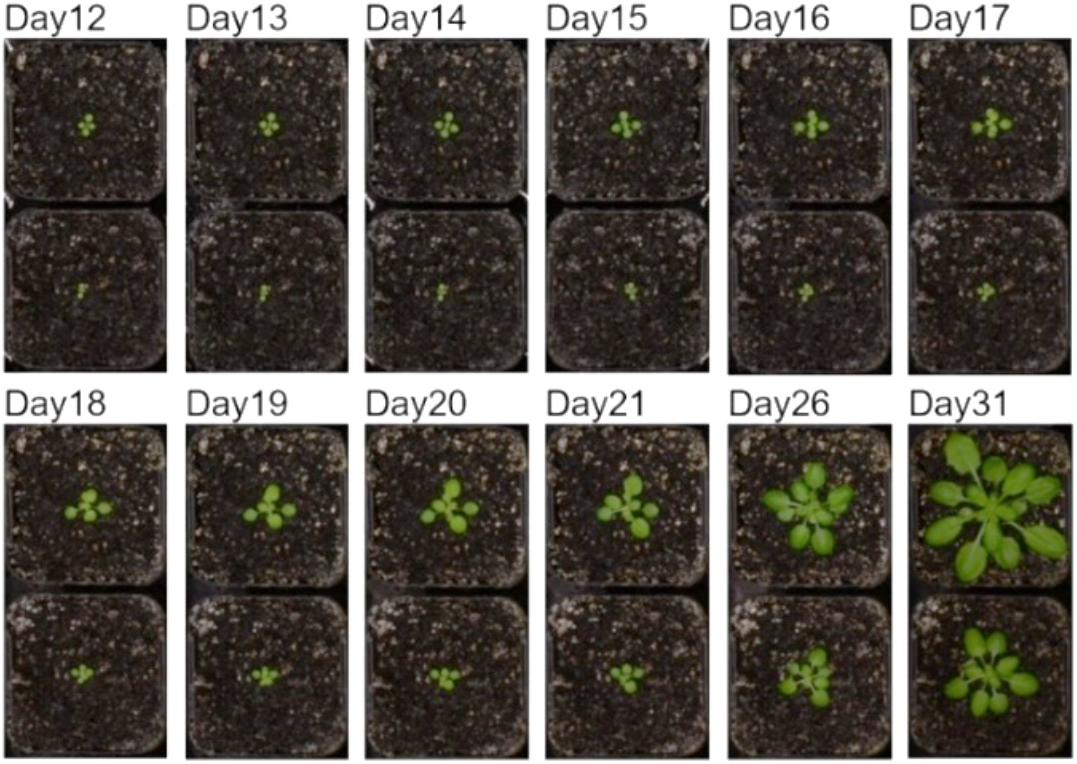
Developmental differences in two neighboring Arabidopsis seedlings (ecotype Col-0) grown under uniform environmental conditions.

The rosette area measurements of all individual Col-0 and Cvi plants were obtained at different time points during the daytime from 12 to 32 days. The mean and standard deviation of these non-normalized measurements were calculated (Figure 2). The overall dispersion of the non-normalized datasets consistently grew from day 12 to day 32, as assessed from the standard deviation of the rosette area measurements (Figure 2 a-c).

**Figure 2.**
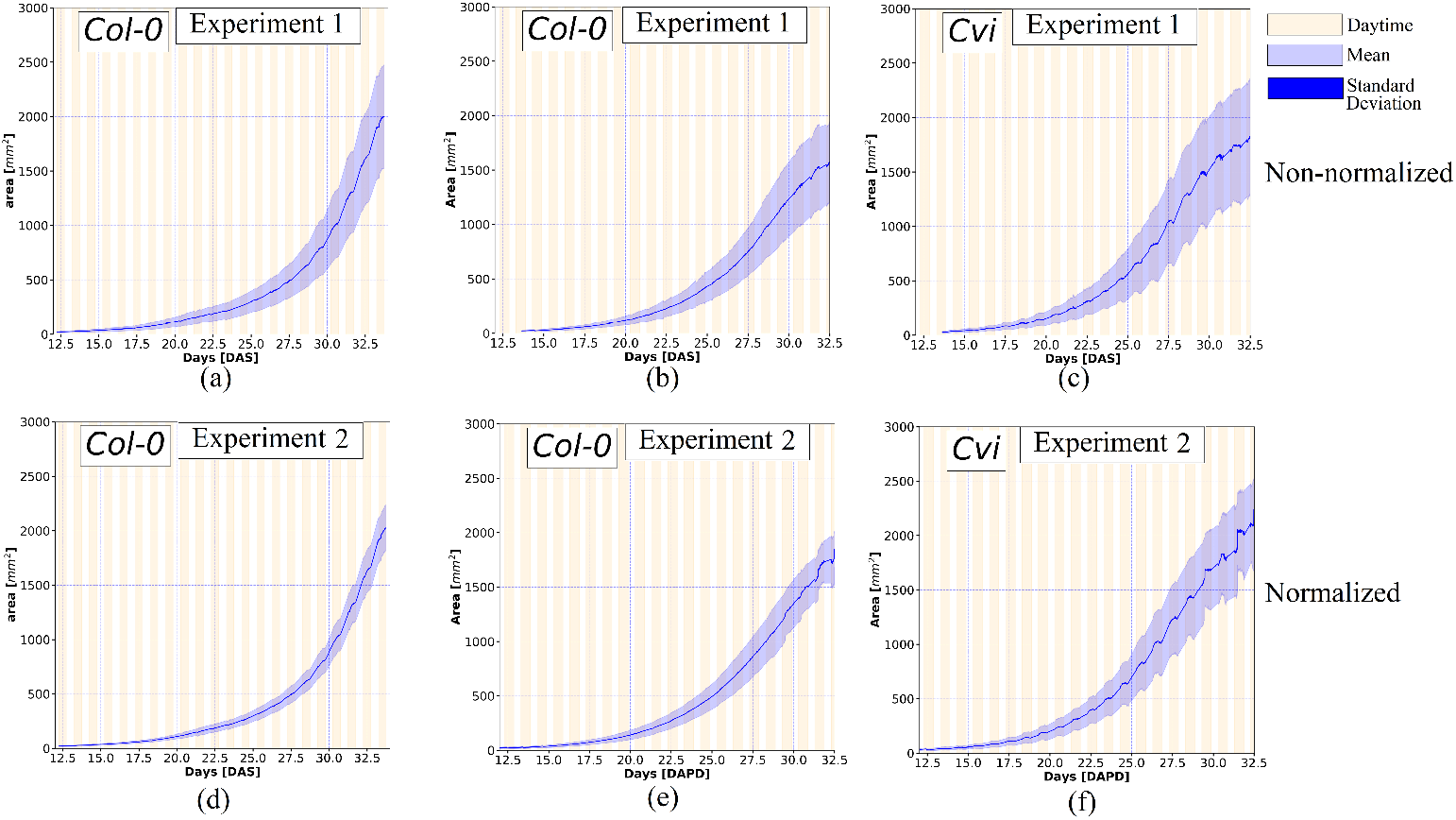
Mean and standard deviation of the non-normalized and normalized projected rosette area datasets. The top row shows the non-normalized datasets: (a) Col-0 plants in experiment 1, (b) Col-0 plants in experiment 2, and (c) Cvi plants in experiment 2. The bottom row shows the normalized datasets: (d) Col-0 plants in experiment 1, (e) Col-0 plants in experiment 2, and (f) Cvi plants in experiment 2. The light purple strip indicates the standard deviation and the solid blue curve in the mean area.

DAPD was applied to normalize the Col-0 and Cvi rosette area datasets. The standard deviation and mean values were calculated at multiple time points to assess the dispersion of the data (Figure 2d-f). The normalized data preserved the exponential growth pattern observed in the non-normalized data, but the overall statistical dispersion of the normalized data was considerably smaller. Daily oscillations in leaf area were also observed, most notably in the Cvi dataset. These occur due to the diurnal change in elevation angle of Arabidopsis leaves, which increases and decreases the angle of the leaves relative to the cameras. Notably, DAPD preserves these signals post-normalization because it shifts the time-series in whole day increments.

Examining the standard deviation over the time-series confirmed the observation that DAPD time normalization reduces the dispersion of rosette area measurments (Figure 3). During the period from day 13 to day 32, the standard deviation of the non-normalized dataset exponentially increased in the three datasets whilst the standard deviation of the normalized dataset linearly increased. DAPD normalization reduced the dispersion of the measurements at day 32 by between 1.5 and 3.5 times.

**Figure 3.**
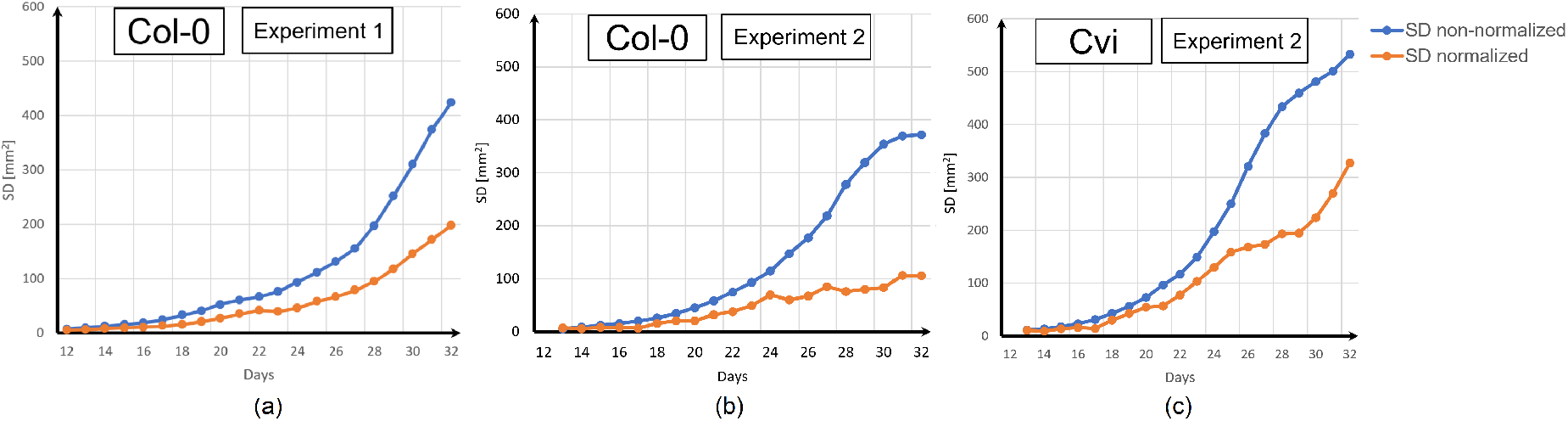
Standard deviation (SD) comparison between the non-normalized and normalized datasets. (a) Col-0 plants in experiment 1, (b) Col-0 plants in experiment 2, and (c) Cvi plants experiment 2.

### DAPD detects outliers with different growth traits amongst a population

A powerful application of DAPD is detection of outliers in HTPP datasets. It is common in large-scale phenomic experiments to observe a small number of individual plants that develop abnormally, despite being of the same genotype as all other members of the population and being grown in uniform conditions. These individuals may have been affected by unintended stresses, and, in some situations, it is reasonable to remove them from datasets. We assessed the ability of DAPD to detect outliers on 24 wild-type Col-0 individuals grown two different trays (Figure 4). It was difficult to confidently identify outliers amongst the non-normalized growth curves by visual inspection or application of a central tendency metric (mean, median, mode of rosette area). However, after applying DAPD normalization, a putative outlier was identified clearly. This plant did not follow the same growth trajectory as the rest of the population, having a smaller rosette area from day 20 onwards. Visual inspection determined that this plant was infected by a pathogen (Figure 5). These results demonstrate DAPD can be applied to detect outliers in HTPP datasets systematically.

**Figure 4.**
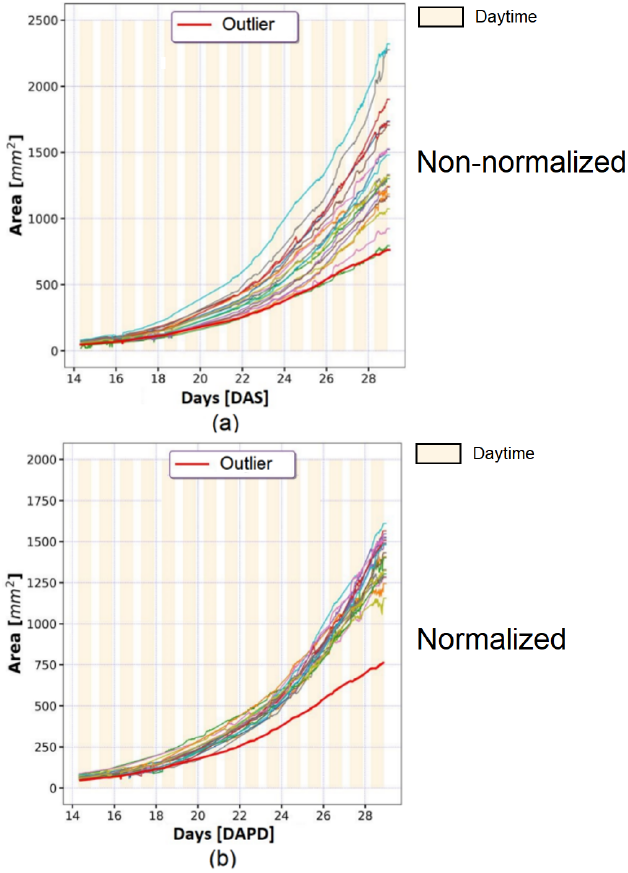
Non-normalized and normalized rosette area curves of 24 Col-0 individuals in experiment 1, (a) non-normalized Col-0 curves, and (b) normalized Col-0 curves. The red and thick curve represents the rosette area of a pathogen-infected individual (outlier) before and after normalization.

**Figure 5.**
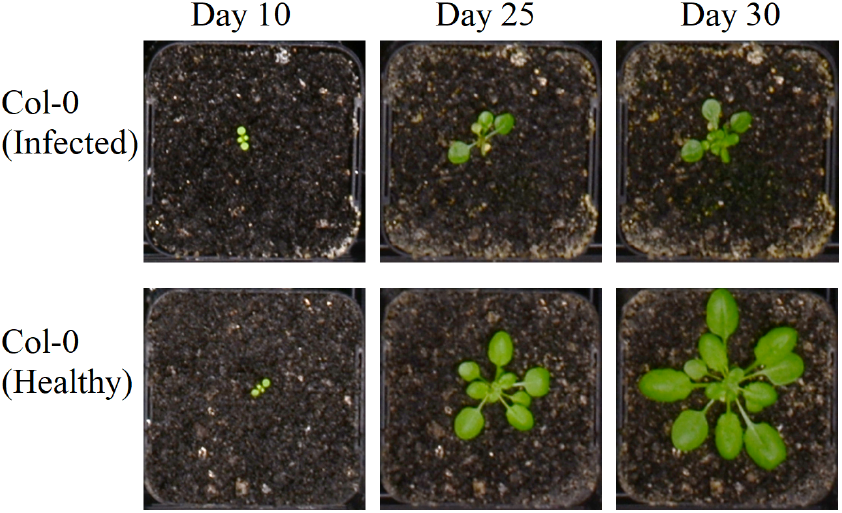
A pathogen-infected plant with outlier growth traits detected automatically using DAPD. The pathogen infected plant had a smaller number of leaves and rosette area than the healthy plant.

### DAPD image segmentation out-performs other image segmentation methods

DAPD depends upon a novel image segmentation method that we developed to improve accuracy and applicability across new datasets compared with existing methods. The method depends on an algorithm that combines shape analysis and supervised machine learning. We benchmarked the accuracy of the DAPD image segmentation algorithm on public datasets (A1, A2, A3, A4) and our in-house generated dataset using dice, precision, recall, and Jaccard metrics (Table 1) [11], [12]. These datasets contain ground-truth RGB and binary images of *A. thaliana* and tobacco plants. DAPD performed consistently well in all metrics across all five datasets. Notably, performance on the tobacco images (A3) comparable to performance on the *A. thaliana* images, indicating DAPD is adaptable to plants with different structure.

**Table 1.**
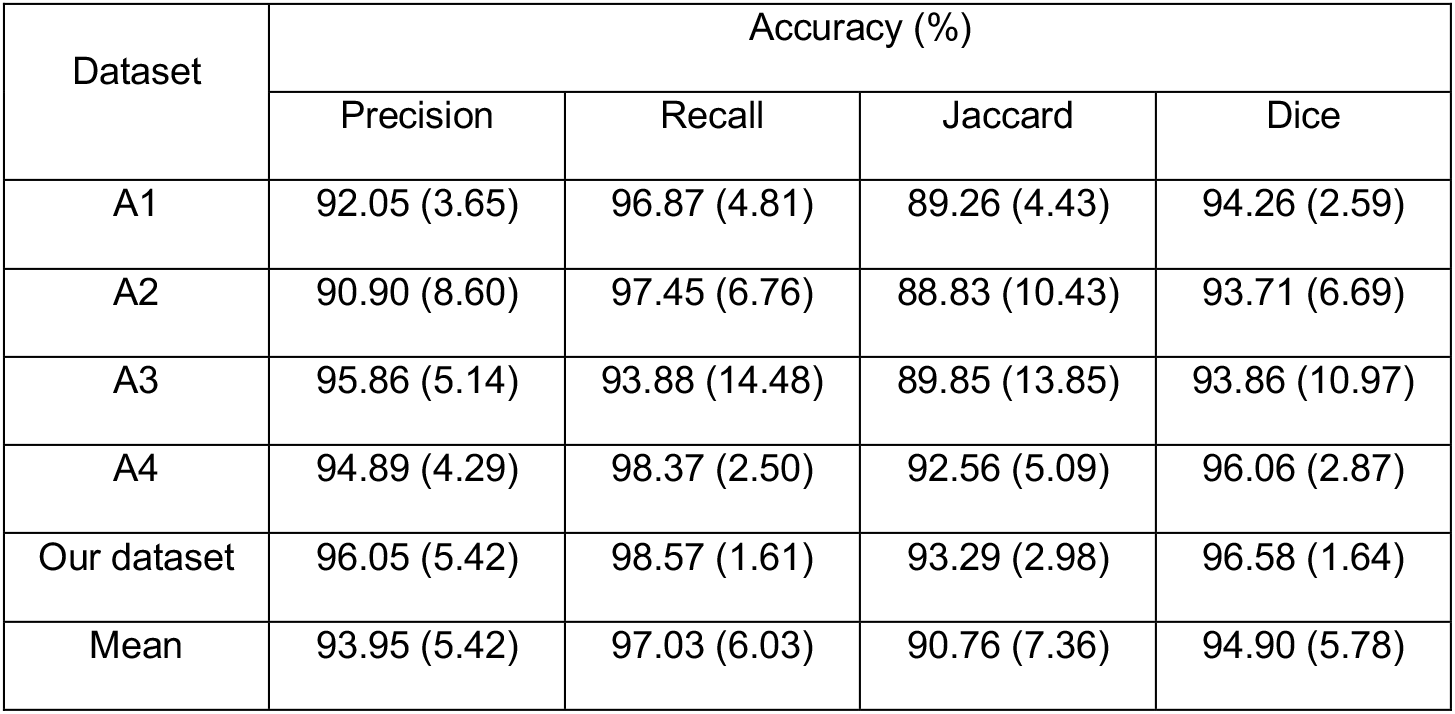
Accuracy results of the DAPD image segmentation algorithm on four public datasets and our dataset. The numerical values in each cell represent the mean and standard deviation (in parentheses). The last row indicates the overall performance of our algorithm.

| Dataset | Accuracy (%) |  |  |  |
| --- | --- | --- | --- | --- |
|  | Precision | Recall | Jaccard | Dice |
| A1 | 92.05 (3.65) | 96.87 (4.81) | 89.26 (4.43) | 94.26 (2.59) |
| A2 | 90.90 (8.60) | 97.45 (6.76) | 88.83 (10.43) | 93.71 (6.69) |
| A3 | 95.86 (5.14) | 93.88 (14.48) | 89.85 (13.85) | 93.86 (10.97) |
| A4 | 94.89 (4.29) | 98.37 (2.50) | 92.56 (5.09) | 96.06 (2.87) |
| Our dataset | 96.05 (5.42) | 98.57 (1.61) | 93.29 (2.98) | 96.58 (1.64) |
| Mean | 93.95 (5.42) | 97.03 (6.03) | 90.76 (7.36) | 94.90 (5.78) |

We compared the performance of the DAPD image segmentation algorithm with three others (Table 2); the Rosette Tracker algorithm [13], the probabilistic parametric active contours algorithm [14], and the Image-based plant phenotyping with incremental learning and active contours [11]. The comparison was conducted by running all five algorithms on the same datasets (A1, A2, A3, A4, and our dataset) and calculating the mean of each performance metric. DAPD image segmentation strongly out-performed active contours and rosette tracker in three of four metrics. Incremental learning performed well but was still not as strong as DAPD image segmentation. These results demonstrated DAPD image segmentation performs well relative to other commonly used approaches.

**Table 2.** Accuracy comparison between active contours, rosette tracker, incremental learning, graph segmentation, and DADP algorithm. Each cell shows the mean and standard deviation of the four metrics (precision, recall, Jaccard, and dice) across the five datasets (A1, A2, A3, A4, and our dataset).

| Method / Algorithm | Accuracy (%)<br>(A1, A2, A3, A4 and our dataset) |  |  |  |
| --- | --- | --- | --- | --- |
|  | Precision | Recall | Jaccard | Dice |
| Active contours | 42.25 (26.05) | 99.66 (26.18) | 42.19 (25.66) | 59.34 (26.20) |
| Rosette Tracker | 54.29 (18.11) | 99.97 (17.79) | 54.29 (19.22) | 70.37 (15.00) |
| Incremental learning | 89.87 (13.68) | 91.94 (2.96) | 83.90 (14.44) | 89.72 (12.36) |
| DAPD | 93.95 (5.42) | 97.03 (6.03) | 90.76 (7.36) | 94.90 (5.78) |

## Discussion

Traditional growth analysis in HTPP systems relies on sowing-time as the temporal start point. This assumption could lead researchers to believe, for instance, that application of an experimental treatment elicits changes in growth at a given time point. However, these analyses may be confounded by differences in germination time or true leaf emergence between genotypes.

We propose the DAPD method to control for temporal differences in development within plant phenotyping datasets. This method uses plant developmental stages to normalize the timeline of phenotyping measurements.

We demonstrate the utility of DAPD to normalize rosette area measurements, but it can be similarly applied to normalize any phenotypic measurements. Furthermore, the analysis we present is on Arabidopsis plants, but DAPD could be used to normalize HTPP data from any species, with two contingencies: First, that a defined and relevant developmental scale can be provided to normalize to, and; second, that an algorithm is available to measure the phenotypic feature of interest in high throughput.

DAPD normalization improved the detection of outliers. Typically a small proportion of plants develop abnormally in large-scale experiments. There are many causes, such as seed quality, low-frequency pathogen infection or unintended stress. The DAPD method enabled clear, systematic detection of anomalous individuals (outliers) within datasets, which is not possible by inspecting central tendency metrics of the non-normalized datasets. DAPD includes a new algorithm to extract the rosette area and count the number of leaves from top-view images. This method outperformed other image segmentation methods when accuracy metrics were assessed on five different ground-truth datasets.

Comparisons between traditional analysis, where sowing time is taken as the reference point, with DAPD normalization illustrated that DAPD decreased dispersion of measurements. The difference in data dispersion between the two approaches gradually increased from the start of experiments to the end. This occurred because dispersion increased more rapidly over the plant lifecycle in the traditional (non-normalized) analysis than the DAPD normalized data. In principle DAPD normalization will enable more sensitive detection of significant differences in trait values between plant lines or ecotypes, due to this decreased variance in measurements. DAPD normalization might also identify differences in germination phenotypes that might otherwise go unnoticed. For example, a mutant with an apparently greater rosette area than wild type during an early growth stage might either have a higher growth rate or germinate earlier. Assuming development was otherwise unchanged, DAPD normalization would eliminate the former possibility, allowing researchers to focus on subsequent experiments. Our code is available for reuse at https://github.com/diloc/DAPD_Normalization.git.

## Methods

### Plant material and growth conditions

Seeds were sterilized in a desiccator using chlorine gas for 150 minutes then stratified for three days in 0.1% agarose in the dark at 4°C. Afterwards the seeds were sown in vermiculite, perlite, and soil mixture (1:1:3), with 20 pots per tray. After germination, only a single seedling was retained per pot. The seedlings were grown in a controlled environment room at 20°C with 50% humidity and were watered with 500 ml water every four days after sowing. Illumination was with an LED light source; it used seven light wavelengths including near-infrared (850 ηm), far-red (740 ηm), deep red (660 ηm), red (618-630 ηm), green (530 ηm), blue (450-460 ηm) and cold white, supplied by PSI Instruments. The average irradiance output on the chamber was set at 150μmol m^-2^ s^-1^ in the photosynthetically active radiation spectrum.

In experiment 1, three hundred and fifty-five individual plants of *A. thaliana* wild-type (Col-0) were grown under 12 h light / 12h dark cycle. In experiment 2, one hundred and forty individuals each of *A. thaliana* ecotypes Cvi and Col-0 were grown under long-day conditions (14 h of light/10 h of dark).

### Imaging

Images were acquired every 15 minutes during the daytime using an HTPP system with 30 RGB cameras that formed a stereo vision system. After the acquisition, images were pre-processed to reduce the noise and correct the color and lens distortion. Subsequently, image segmentation algorithms extracted the rosette area and shoot phenotypic measurements such as the rosette area, growth rate, and leaf number were calculated. The image processing steps and details are described in the following.

### Image pre-processing

Image pre-processing algorithms corrected the lens and color distortion. The lens correction algorithm calculated the optimal rotation and translation camera parameters using a chessboard pattern and the pin-hole camera model. Color distortions were removed using a mapping function between the light intensity and color checker cards, which had been included in the imaged area.

### Rosette segmentation

The projected rosette area of individual plants was extracted from pre-processed images by using a combination of multiple image segmentation algorithms. First, individual pot regions were dynamically cropped from the RGB images using an adaptive window. The resultant image was converted to HSV color space and the value (V) channel extracted from it. The V channel was used to enhance the contrast of the image by applying the Contrast Limited Adaptive Histogram Equalization (CLAHE) technique [15]. Subsequently, the green color component of the image was obtained by the independent contribution of the RGB and HSV color space. The green component image was binarized using Otsu’s method for global automatic thresholding [16]. The salt-and-pepper noise of the Otsu result was removed by applying a median filter, followed by using a Gaussian filter, which reduced high-frequency spatial noise. The binary image was further processed by an area-fill operation to remove small unwanted background regions or holes.

The resultant binary image was used to segment the plant shoot from the original HSV image. Undesired background objects were identified and removed from it by grouping into coherent classes using K-means clustering [17]. Then, a histogram of the Hue channel was extracted and smoothed using the Savitzky-Golay filter [18]. A complementary spatial correlation test was used to determine the rosette objects. The test checked the correlation between image clusters. These clusters were formed by selecting all colors in a distribution except by color values beyond the curve’s intersection point. Spatially correlated clusters belonged to the same object and were retained. Non-correlated clusters were removed. The whole procedure was repeated multiple times, and at each time, the gamma value was randomly modified. This iterative approach occasionally over-segmented some rosette areas, but it was corrected by using Kriging [19]. Kriging is also known as Gaussian process regression, which produced predictions of unobserved values from observations at nearby locations [20]. Finally, the projected rosette area was obtained from the segmented rosette image. Unexpected power disruptions caused some image data not to be acquired during the time-series. Considering the complete time series as being images acquired every 15 minutes during the day, the data missing from experiment 1 was 14.83% of the total data and in experiment 2 was 5.6%. To enable consistent downstream analyses we applied data imputation techniques to estimate the missing data, including up-sampling to a higher frequency, spline interpolation and curve fitting.

### Individual leaf Segmentation / Counting (DAPD segmentation)

Our leaf segmentation approach combined two algorithms; edge contour extraction and marker-controlled watershed [21]. The edge contour extraction process was applied to the rosette binary image to obtain the rosette contour using topological analysis [22]. The center of mass of the rosette was calculated using the Hough transformation. This information was used to calculate leaf morphology features such as tip, base, center, and petiole. The watershed transformation was applied to the segmented rosette image, and centers of leaves were used as initial markers. This identified overlapping leaves and separated convex and smooth rosette features that touched. After applying the watershed transformation, the resulted number of markers represented the total number of leaves in the rosette.

### Developmental normalization (DAPD normalization)

The normalization by development removed the time difference between plant measurements, which had similar developmental stages. These stages were identified by the leaf number based on the adjusted BBCH scale [3].

The leaf number of each plant was measured by applying the DAPD segmentation algorithm to the rosette images. The measured leaf number was a discrete function (1), which depended on the real leaf number, leaf occlusion effect, and modeling error denoted ME(t).

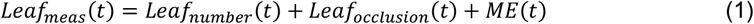

The variability of the measured leaf number function (1) was minimized by calculating the trend using curve fitting. This trend assembled an exponential function in the early developmental stages (2). The coefficient was the initial value of the function, and *b* was the growth rate.

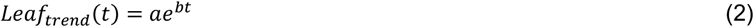

The leaf number trend may vary from plant to plant within the same line/mutant population at a time point. Then, the average leaf number trend among all individuals was calculated to homogenize the trends (3).

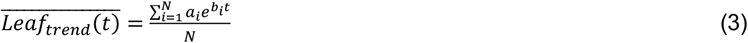

The time-difference of plants with similar leaf number was removed by finding the best timeline in (1) that fitted (3). Multiple timelines were generated from the original timeline (t) by inserting a time delay (k). This time delay could be a positive or negative integer number that shifted the original timeline in days (t – k). The mean squared deviation (MSD) was calculated per each time delay (4). The time delay (s) that produced the lowest MSD was selected to shift the leaf number (5) and rosette area (6) time series. However, this time delay (s) must be adjusted because plants could have the same number of leaves, but the rosette area might be different due to the maturity and expansion of leaves. Δ*t* represented the time delay adjustment, as shown in equation (7).

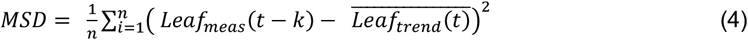

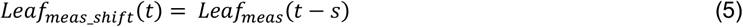

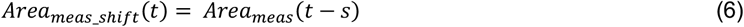

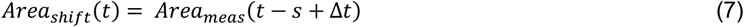

## Abbreviations

BASF: German for Baden Aniline and Soda Factory
BBCH: BASF, Bayer, Ciba-Geigy
CLAHE: Contrast limited adaptive histogram equalization
DAPD: digital adjustment of plant development
DAS: day-after-sowing
HSV: hue, saturation, value
HTPP: high throughput plant phenotyping
LED: light-emitting diode
RGB: red, green, blue

## Authors’ contributions

DL, OB, JW and ML designed the experiments. DL, XM and OB performed the experiments. DL acquired data, processed images and analyzed data. DL, GD and EC developed algorithms. DL, OB, EC, GD and ML interpreted the results and wrote the paper. All authors read and approved the final manuscript.

## Acknowledgments

We thank Andrew Robinson for helping set up the infrastructure for data storage and transfer. We thank Dr. Ricarda Jost and Meiyan Ren for assisting with experiments. We thank Dr. Martin Trtilek and Photon Systems Instrument for building the HTPP system.

## Availability of data and materials

Our code is available for reuse at https://github.com/diloc/DAPD_Normalization.git. Correspondence should be addressed to.

## Consent for publication

All authors reviewed and approved the final version of the manuscript for submission.

## Ethics approval and consent to participate

Not applicable.

